# The structural basis for activation of voltage sensor domains in an ion channel TPC1

**DOI:** 10.1101/373860

**Authors:** Alexander F. Kintzert, Evan M. Green, Pawel K. Dominik, Michael Bridges, Jean-Paul Armache, Dawid Deneka, Sangwoo S. Kim, Wayne Hubbell, Anthony A. Kossiakoff, Yifan Cheng, Robert M. Stroud

**Affiliations:** University of California, San Francisco, Department of Biochemistry and Biophysics; University of Chicago, Department of Biochemistry and Molecular Biology; University of California, Los Angeles, Jules Stein Eye Institute and Department of Chemistry and Biochemistry; Howard Hughes Medical Institute

## Abstract

Voltage sensing domains (VSDs) couple changes in transmembrane electrical potential to conformational changes that regulate ion conductance through a central channel. Positively charged amino acids inside each sensor cooperatively respond to changes in voltage. Our previous structure of a TPC1 channel captured the first example of a resting-state VSD in an intact ion channel. To generate an activated state VSD in the same channel we removed the luminal inhibitory Ca^2+^-binding site (Ca_i_^2+^), that shifts voltage-dependent opening to more negative voltage and activation at 0 mV. Cryo-EM reveals two coexisting structures of the VSD, an intermediate state 1 that partially closes access to the cytoplasmic side, but remains occluded on the luminal side and an intermediate activated state 2 in which the cytoplasmic solvent access to the gating charges closes, while luminal access partially opens. Activation can be thought of as moving a hydrophobic insulating region of the VSD from the external side, to an alternate grouping on the internal side. This effectively moves the gating charges from the inside potential to that of the outside. Activation also requires binding of Ca^2+^ to a cytoplasmic site (Ca_a_^2+^). An X-ray structure with Ca_a_^2+^ removed and a near-atomic resolution cryo-EM structure with Ca_i_^2+^ removed define how dramatic conformational changes in the cytoplasmic domains may communicate with the VSD during activation. Together four structures provide a basis for understanding the voltage dependent transition from resting to activated state, the tuning of VSD by thermodynamic stability, and this channel’s requirement of cytoplasmic Ca^2+^-ions for activation.

## Introduction

Voltage sensing domains (VSDs) are four-helical bundle domains, termed S1-S4 that respond to changes in membrane potential by allowing ‘gating’ charges, generally positively charged arginine or occasionally lysine side chains in the fourth transmembrane helix S4 (charges referred to as R1-R5) to move relative to a charge-transfer center (CT) (1, 2) that contains counter charges in the surrounding helices S1-S3 and an aromatic residue (Y, F) that seals the VSD to solvent passage.

The number of gating charges in each VSD that move across the membrane from connection to the cytoplasmic side to the extracellular (or lumenal) side during activation is typically measured as 2-3 and up to 5 positive charges. The basis for structural and electrical changes in S4 that give rise to voltage-dependence is key to understanding the response of voltage-gated ion channels to changes in membrane potential.

In voltage-gated ion channels, the movement of S4 is connected via an S4-S5 linker helix on the cytoplasmic side to the pore helices S5-S6 of the ion channel, and also through Van der Waals hydrophobic contacts between S4 and the pore-forming domains. In excitable cells, activation of the VSD in response to membrane depolarization greatly increases the probability of channel opening(3, 4).

Several models have been proposed for voltage-dependent activation. The ‘Sliding Helix’(5), ‘Rotating helix’(6, 7), and ‘Paddle’(8) models suggest that S4 moves substantially (vertically or in rotation) in the membrane to translate the gating charges across the membrane (Figure S1). In Sliding Helix and Rotating Helix models, the gating charges interact with counter anions or aqueous environments to avoid the energetic penalty of placing a charge in a hydrophobic environment. Lanthanide-based resonance energy transfer measurements suggest that gating charges do not move extensively during activation, but rather achieve alternating exposure to the internal and external milieu through conformational changes in the VSD helices S1-S4 (9). A ‘ratchet’ model also includes the possibility of multiple intermediate states of S4(10). Electrophysiology, electron paramagnetic resonance (EPR) spectroscopy, X-ray structures, disulfide crosslinking, and simulations support a combination of translation and rotation of S4 during activation (11–18). No structural information until now exists for voltage-gated channels captured in multiple activation states, precluding an atomic scale evaluation of the mechanism of voltage-dependent changes, or how they translate to channel activation.

Our previous crystal structure of wild-type *Arabidopsis thaliana* two-pore channel 1 (AtTPC1_WT_) provided the first resting-closed state in an intact ion channel(19, 20), the electrophysiological state that forms under high lumenal 1 mM (>100μM) Ca^2+^-ion concentration. Here we sought to determine structures for the activated-state of AtTPC1 that is formed in low lumenal (<100μM) Ca^2+^-ions by removing the inhibitory luminal Ca^2+^-binding site in VSD2 (Ca_i_^2+^), while keeping the cytoplasmic activation site (Ca_a_^2+^) occupied with >300μM cytoplasmic Ca^2+^-ions (21). Secondly, we wanted to determine the mechanism for the channel’s requirement of cytoplasmic Ca^2+^-ions for activation by removing the Ca_a_^2+^ site.

TPCs are a family of ion channels that regulate ion conductance across endolysosomal membranes(21, 22, 23). Located in endosomes that endocytose from the plasma membrane, initially with ~1mM extracellular Ca^2+^ concentration, they regulate the conductance of Na^+^-and/or Ca^2+^-ions out of the endolysosome, intravesicular pH (24), trafficking (25), and membrane excitability (26). Cytoplasmic Ca^2+^-ions (>300 μM) are required for any activation of AtTPC1 (27), whereas lumenal Ca^2+^-ions (>100 μM) suppress voltage-dependent activation (20, 28). TPCs encode two pore-forming domains on a single chain with two non-equivalent VSDs (S1-S4, S7-S10) and pore helices (S5-S6, S11-S12). In AtTPC1 only VSD2s (S7-S10) respond to changes in voltage (20). Three arginines on S10 of each VSD2 (equivalent to S4 in the VSD of tetrameric ion channels) in AtTPC1 are required for voltage-dependent activation. A homo-dimer of two TPCs forms the central functional channel surrounded by four pore-forming domains. The dependence of AtTPC1 on external and internal Ca^2+^ offer the opportunity to visualize the resting state of a voltage sensing domain of an intact channel, and the activated state and to ask how voltage changes are detected and relayed.

Lumenal Ca^2+^-ions suppress activation of AtTPC1 via binding to Ca_i_^2+^ located in the active voltage sensor VSD2 with EC_50_ ~0.1mM. This previously enabled trapping of the resting-state VSD2 of wild-type AtTPC1 by including 1 mM Ca^2+^-ions (19, 20). Replacement of the Ca^2+^-chelating amino acids by mutagenesis (D240N, D454N, E528Q; termed AtTPC1_DDE_) shifts voltage-dependent activation by −50 mV, such that the channel is open at 0 mV (20, 28) and VSD2 in an activated conformation. Using AtTPC1_DDE_ we sought to determine the structure of an activated state of the same intact channel where we had previously determined a resting state.

Channel opening requires Ca^2+^-ion binding to Ca_a_^2+^ mediated by D376 of cytoplasmic EF-hand domain helix 3-4 loop (EF3-EF4). Removal of cytoplasmic Ca^2+^-ions or the mutation D376A (AtTPC1_DA_) yields permanently closed channels (29). The absolute requirement of the Ca_a_^2+^ site for voltage-dependent activation led us to hypothesize that channel activation depends on communication between Ca_a_^2+^ and VSD2 (21). Using the D376A mutation we sought to determine how cytoplasmic Ca^2+^ evokes activation.

## Results and Discussion

### Cryo-EM structures of AtTPC1_DDE_

As a basis for understanding the voltage-dependent activation mechanism and its modulation by Ca^2+^-ions, we determined the structure of AtTPC1_DDE_ by cryo-EM (Figure 1a, Figure S2, S3, S4, Table S1). We employed saposin A nanoparticles (30) to reconstitute AtTPC1_DDE_ into a membrane environment, and an antibody Fab made against AtTPC1 (CAT06/H12) (31) to facilitate particle alignment (32) (Figures S5, S6, See Supplemental Discussion).

**Figure 1.**
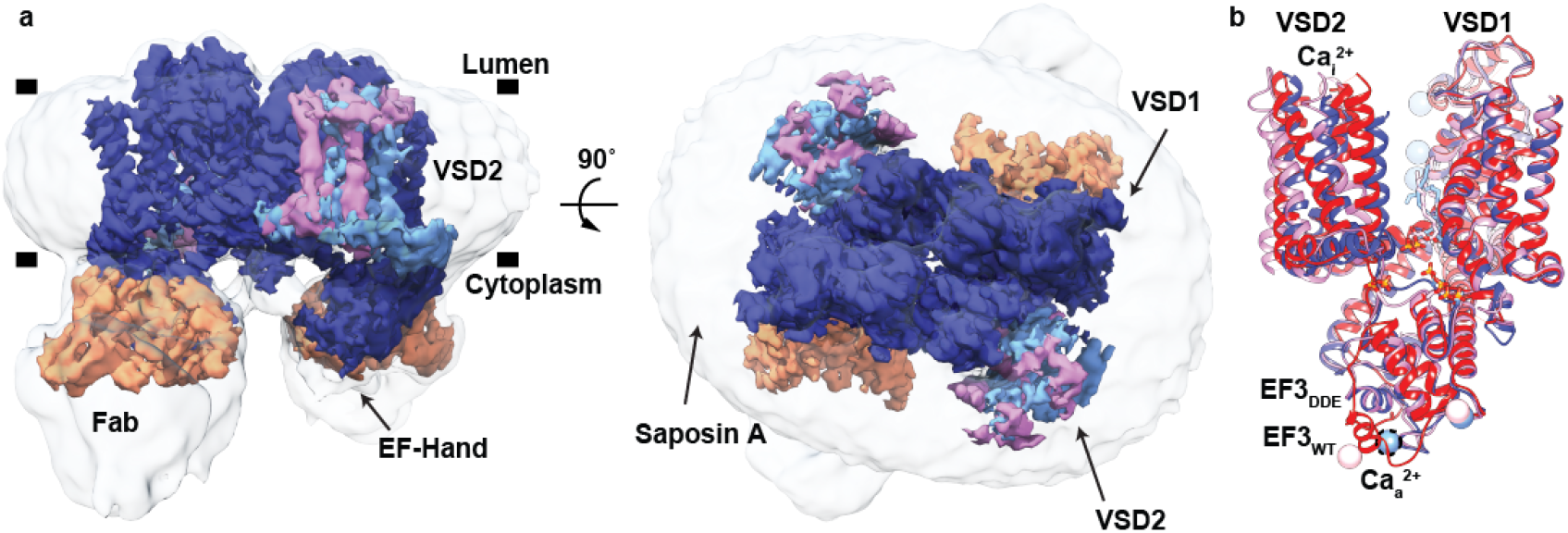
Cryo-EM Structure of the AtTPC1_DDE_-saposin A-Fab complex. a, Side (left) and top-down (right) views of cryo-EM density. The composite map is colored to highlight high-resolution features of TPC1 (blue, EMDB 8957), VSD2 in state 1 (cyan, EMDB 8958) and state 2 (pink, EMDB 8960), and the Fab variable domains (orange, EMDB 8956). Unsharpened density is shown at low contour (transparent gray). Membrane boundaries (black bars) defined by the nanodisc are marked. b, Overlay of AtTPC1_WT_ (red, PDBID 5DQQ), state 1 (blue), and state 2 (pink). Ca_i_^2+^ and Ca_a_^2+^ denote sites of lumenal inhibition and cytoplasmic activation by Ca^2+^-ions. Dash lines around Ca_a_^2+^ indicate hypothetical position in cryo-EM structure. Ca^2+^-ions shown as colored balls.

With a nominal resolution of 3.3 Å, the density map of AtTPC1_DDE_ is of high quality (Figure S4a), allowing additional *de-novo* interpretation of the AtTPC1 N-terminal domain (NTD), the S1-S2 linker, EF3-EF4 with an intact Ca_a_^2+^ site, the upper vestibule of the pore, and the C-terminal domain (CTD). Three ions lie in the selectivity filter, consistent with previously defined Ca^2+^/Na^+^-binding sites (19, 20, 33). Fourteen lipid molecules surround the channel in the lumenal leaflet with two on the cytoplasmic side (Figure S7). S1-S6 are well defined and remain stationary between all of the AtTPC1 structures. In AtTPC1_DDE_, major rearrangements are observed in the VSD2, the upper vestibule of pore, and EF3-EF4 on the cytoplasmic side, relative to AtTPC1_WT_ (Figure 1b). The position of three residues known to be phosphorylated (S22, T26, and T29) are observed in previously unresolved portions of the NTD (Figure S4a, See Supplemental Discussion) (19).

The Fab binds to the EF1-EF2 loop, EF4, and S6 on the cytoplasmic side. The Fab binding affinity is the same with and without Ca^2+^ ions, in other molecular conditions tested (1 mM EGTA, 1 μM trans-NED19, 1 μM Nicotinic Acid Adenine Dinucleotide Phosphate), and in three different constructs (AtTPC1_WT_, AtTPC1_DDE_, AtTPC1_DA_), indicating that the Fab binding does not distinguish between nor is likely to influence the activation state (Figure S5c-d). Most of the variable domain of the Fab is visible, allowing interpretation of the AtTPC1_DDE_-Fab interface (Figure S6e-f). The constant domain of the Fab is flexible and not resolved to high resolution.

### Activation of the voltage sensor

Lumenal Ca^2+^-ions inhibit AtTPC1 channel activation half maximally at ~0.1 mM concentration(20) via binding to Ca_i_^2+^ (21) between VSD2 (D454 in the S7-S8 loop, and E528 in S10), and the pore (D240). D454N, also named *fou2* (fatty acid oxygenation up-regulated 2) (28), abolishes inhibition by lumenal Ca^2+^-ions, increasing channel open probability by shifting the voltage-dependent channel opening towards more hyperpolarizing potentials (28, 20). Ca^2+^-ions do not inhibit AtTPC1_DDE_ at 1 mM and up to 10 mM (33). Thus, the Ca_i_^2+^ site is functionally abolished in AtTPC1_DDE_ under cryo-EM conditions and thereby mimics the activated state under voltage and low lumenal Ca^2+^-ion conditions (20, 33).

While the overall resolution of AtTPC1_DDE_ is excellent, density for S7-S10 of VSD2 is significantly weaker indicating conformational heterogeneity. Focused classification identified two states of VSD2, each with an overall resolution of 3.7 Å, but with distinct and different conformations of the S7-S10 domain structure (state 1 and 2) (Table S1, Figure S3, See Methods). We therefore conclude that these two conformations represent different functional states of the AtTPC1_DDE_ channel. They suggest a model for the voltage-activation of AtTPC1, and a mechanism for dependence on lumenal Ca^2+^-ions.

To confirm that the structural rearrangements in VSD2 were not induced by saposin A we determined a ~7Å reconstruction of AtTPC1_WT_ in both detergent and in saposin A. In these the overall channel architecture is comparable to the AtTPC1_WT_ crystal structure (Figure S8).

The local resolution of VSD2 (S7-S10) ranges from 4-6 Å in both states, making it possible to model the intermediate active state of VSD2 based on the 2.8 Å X-ray structure of AtTPC1_WT_ (Figure S4b, c, See Methods under Structure Determination and Refinement)(19). Atomic structures for states 1 and 2 were determined by real-space and B-factor refinement against the cryo-EM densities. Changes in solvent accessibility at the luminal and cytoplasmic boundaries of VSD2 were apparent from the comparison of these new structures of AtTPC1 and the previous resting-state. Taking the resting state structure of AtTPC1_WT_ as a reference (PDBID 5DDQ; ref. (19)), transmembrane helices S7-S10 in state 1 rotate in a counterclockwise manner with respect to AtTPC1_WT_ (Figure 2). The helices twist to partially close the cytoplasmic solvent access to the gating charges of VSD2 (Figure 3), while the lumenal face remains occluded. Among all helices of VSD2, the key arginine-rich S10 has the least movement, and R537 (R1) remains interacting with the CT (Y475), albeit the changes in S8 conformation place R1 on the opposite horizontal face of the CT as compared with AtTPC1_WT_. S8 moves upward in the membrane plane by nearly one helical turn, thus moving the CT into place to interact with R1 in similar manner to AtTPC1_WT_. State 1 probably represents a resting-state structure present in low lumenal (<0.1 mM) Ca^2+^-ion concentrations.

**Figure 2.**
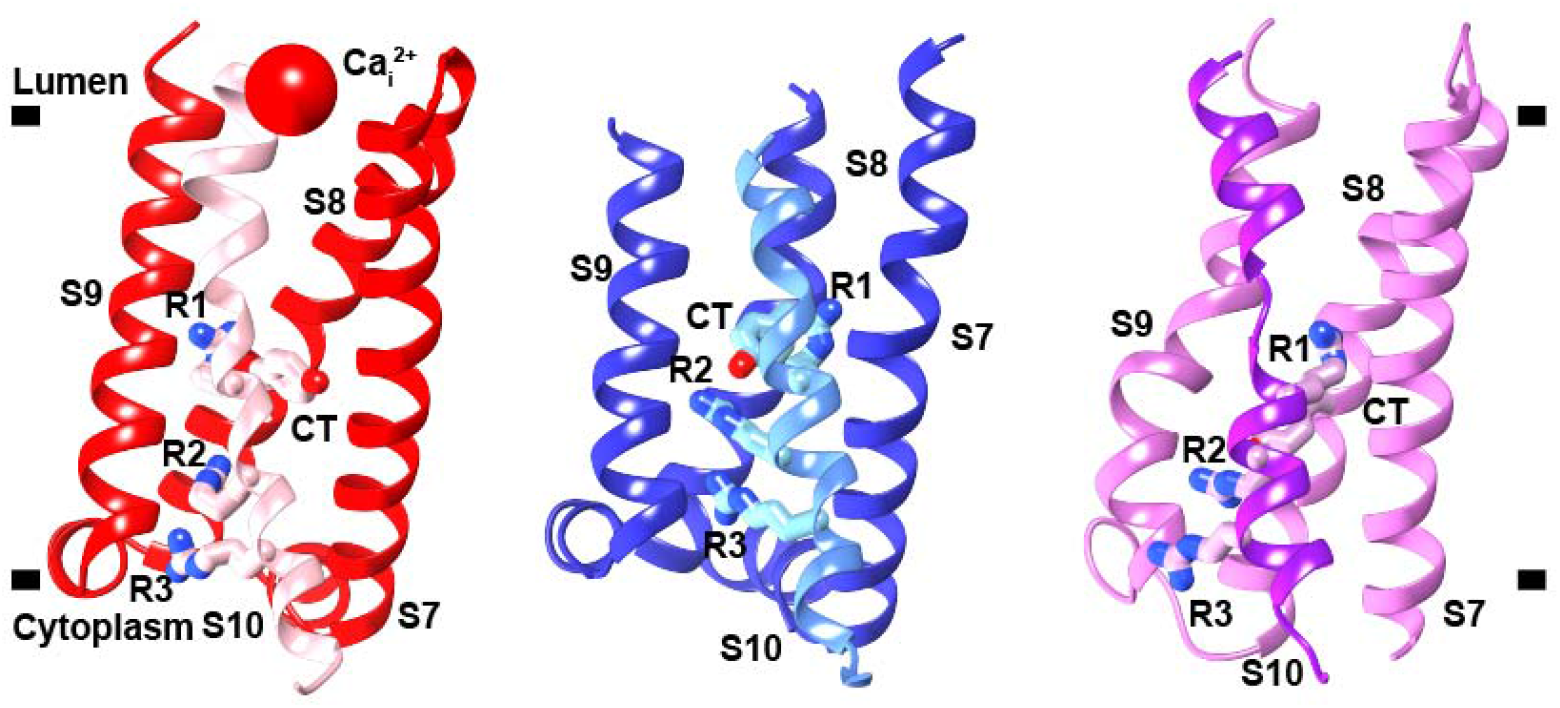
Three States of VSD2. View of VSD2 in the membrane plane from the center of the channel looking outward, that illustrates the rotation and twisting of VSD2 helices S7, S8, S9, and S10. Connections to the pore domains omitted for clarity. S10 is highlighted in each state with a different color than the other helices. Gating charges R1-R3 (R537, R540, R543), and CT residue Y475 are shown. Left: resting-state AtTPC1_WT_ (red, PDBID 5DQQ), Center: AtTPC1_DDE_ state 1 (blue), and Right: AtTPC1_DDE_ state 2 (pink).

In state 2, VSD2 rotates ~20° clockwise in the plane of the membrane with respect to AtTPC1_WT_ (Figure 2). Helices S7-S10 reorient dramatically leading to an opening of the lumenal face of VSD2 (Figure 3). Tilting of S10 and rotation of S8 around S10 moves the CT downward, placing R1 in an activated conformation. The cytoplasmic face is fully closed in state 2 while the lumenal side is partially open. VSD2 has effectively alternated solvent access, or electrical contact from the cytoplasmic to the lumenal side of the membrane.

**Figure 3.**
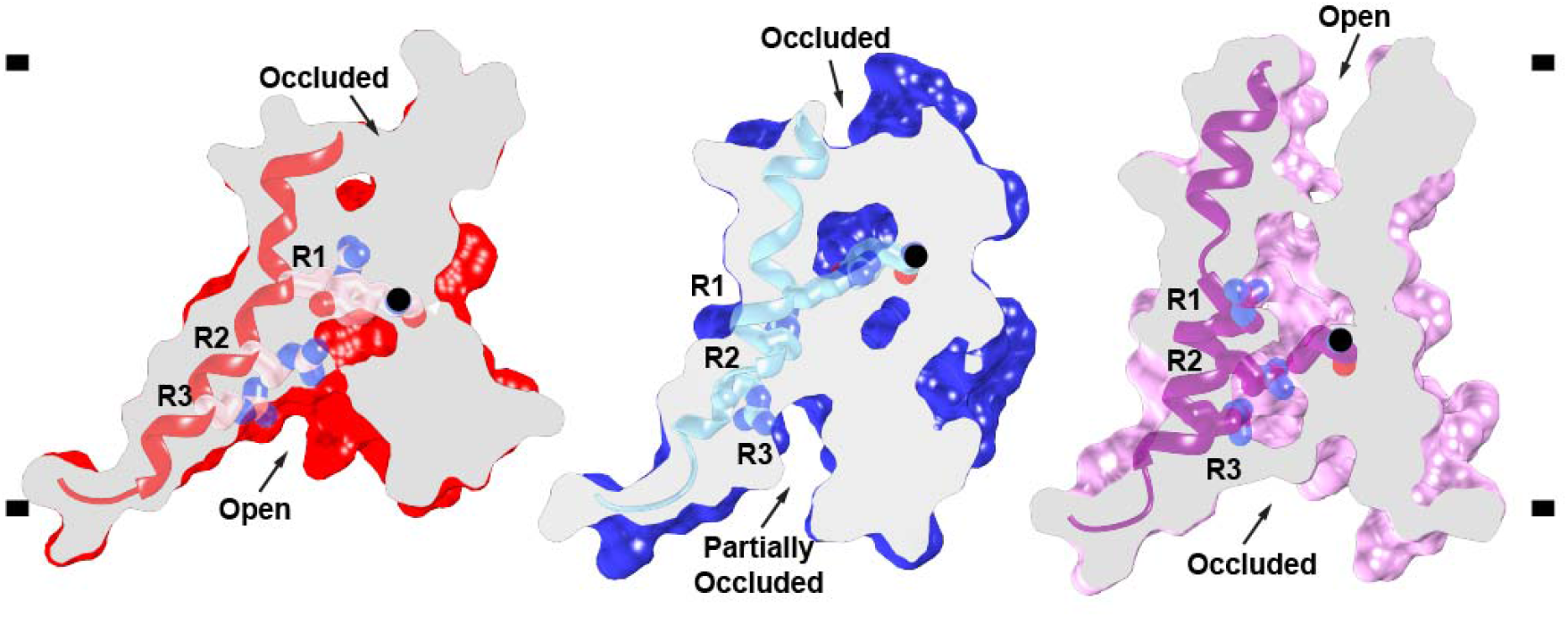
Activation of the Voltage-sensor. Side views of a common slice perpendicular to the membrane surface of VSD2 in AtTPC1_WT_ resting-state (red, PDBID 5DQQ) and AtTPC1_DDE_ state 1 (blue) and state 2 (pink). S10 shown with gating charges R1-R3. The CT C□ position (black ball) is marked. Based on a structural alignment with respect to the pore helices S6-S7 of each structure.

Clockwise rotation was proposed to connect resting- and active-state conformations on the basis of comparing previous voltage-gated channel structures (13, 19, 34). During review of this manuscript, activated state structures of mouse TPC1 were determined (35). Comparing the activated-state mouse TPC1 to resting- and intermediate-state AtTPC1 supports the proposed role of VSD2 rotation during channel activation. The observation of clockwise rotation in multiple activated-state structures supports our conclusion that state 2 represents an intermediate-activated state of AtTPC1.

These two states of VSD2 in AtTPC1 are likely to represent structures that VSD2 adopts in low lumenal Ca^2+^ during activation. The overall movement of VSD2 during activation serves to dilate the lumenal face of VSD2, close off the cytoplasmic leaflet to solvent, and move the CT below a gating charge in the membrane. This would reconnect the positive gating charges from the potential of the cytoplasmic side to the potential of the lumenal side upon activation, mediated by tilt and twisting of S7, S8, S9 around S10. This is all that is necessary to transport the gating charge arginine residues from cytoplasmic electrical connection, to the external potential. A hydrophobic sealed region between all four helices forms a thin outer insulating layer in the resting state, while in the activated state this becomes open to solvent and the bundle twists to form a hydrophobic insulating region closer to the cytoplasmic side.

### Drug and Lipid Binding Sites

The high resolution map enabled refinement of a total of 14 lipids on the lumenal and 2 on the cytoplasmic leaflet of the membrane. The lumenal lipids are modeled as the 16-carbon containing palmitic acid, the predominant lipid length in soy polar lipids, whereas the cytoplasmic lipids are modeled as 18-carbon phosphatidic acid (PA). 12 lipids bind to the long axis of AtTPC1, occupying the binding site for Ned19 (Figure S7, ref. (19, 21)). Ned19 binding may disrupt these structured lipids, acting as a steric block to prevent S7 and pore movements during gating. One lipid binds to a buried site along the short axis, sandwiched in between the pore domains. Based on homology to Cav channels in this region and recent structures of bacterial Cav channels (CavAb) bound to dihydropyridines (DHP) (amlodipine and nimodipine) and PPA (verapamil) inhibitors(36), the short axis in transmembrane segments S6 and S12 is a likely site for binding of Tetrandrine, a bis-benzylisoquinoline alkaloid isolated from the Chinese herb *Stephania* tetrandra(37) and approved medications of the DHP class of L-type Ca_v_ antagonists–all TPC channel blockers.

The lumenal PA lipids bind along the short axis in a pocket formed by S1 of VSD1, the S10-S11 linker, S11, and the S8-S9 linker of VSD2. The alkyl chains make Van der Waals interactions, whereas the phosphoglycerol headgroup makes hydrogen bonds to the backbone of W492 and to the sidechain of R498 and S8-S9 of VSD2. The S8-S9 linker moves 4Å inward in the AtTPC1_DDE_ structure to form the PA binding site, which would not be intact in the AtTPC1_WT_ resting-state. Therefore, this lipid binding site is likely specific for AtTPC1_DDE_. Certain lipids may occupy this site during activation of AtTPC1. Polyunsaturated fatty acids inhibit plant TPC1 activation, but the binding site has not been determined (38). In principle, polyunsaturated fatty acids could mimic the observed site in the resting-state and prevent activation.

### Charge transfer mechanism

State 2 represents an intermediate activation state of VSD2 where the gating charge R1 is transferred across the membrane (Figure 3). In this state, the CT moves downwards, R1 moves slightly upwards, and overall the VSD2 bundle rotates by ~20° in the membrane plane with respect to the adjacent pore domains. In this the voltage-sensing helix S10 pivots only 2-3° inward toward the pore. Importantly, the three observed states of the VSD thus far are not related by rigid-body rotations. These changes in VSD2 make R1 more accessible to solvent on the lumenal side while charges R2 and R3 are shielded from access to the cytoplasmic side; there is a net transfer of charge from cytoplasmic side (in the resting state) to the outside (in the activated state).

### Pore conformational changes

High-resolution density for the pore defines the consequences of activation by removal of the external Ca_i_^2+^ site on VSD2 on the central channel (Figure 4a, b). As a consequence of the rearrangement of VSD2 the upper vestibule and the upper selectivity filter of the channel open along the ion permeation pathway (Figure 4c), while the lower gate remains closed (Figure 4d). Full channel opening may be evoked by passage of ions or an additional energy barrier, because electrophysiological studies show the channel is maximally open at 0 mV with removal of the Ca_i_^2+^ site.

**Figure 4.**
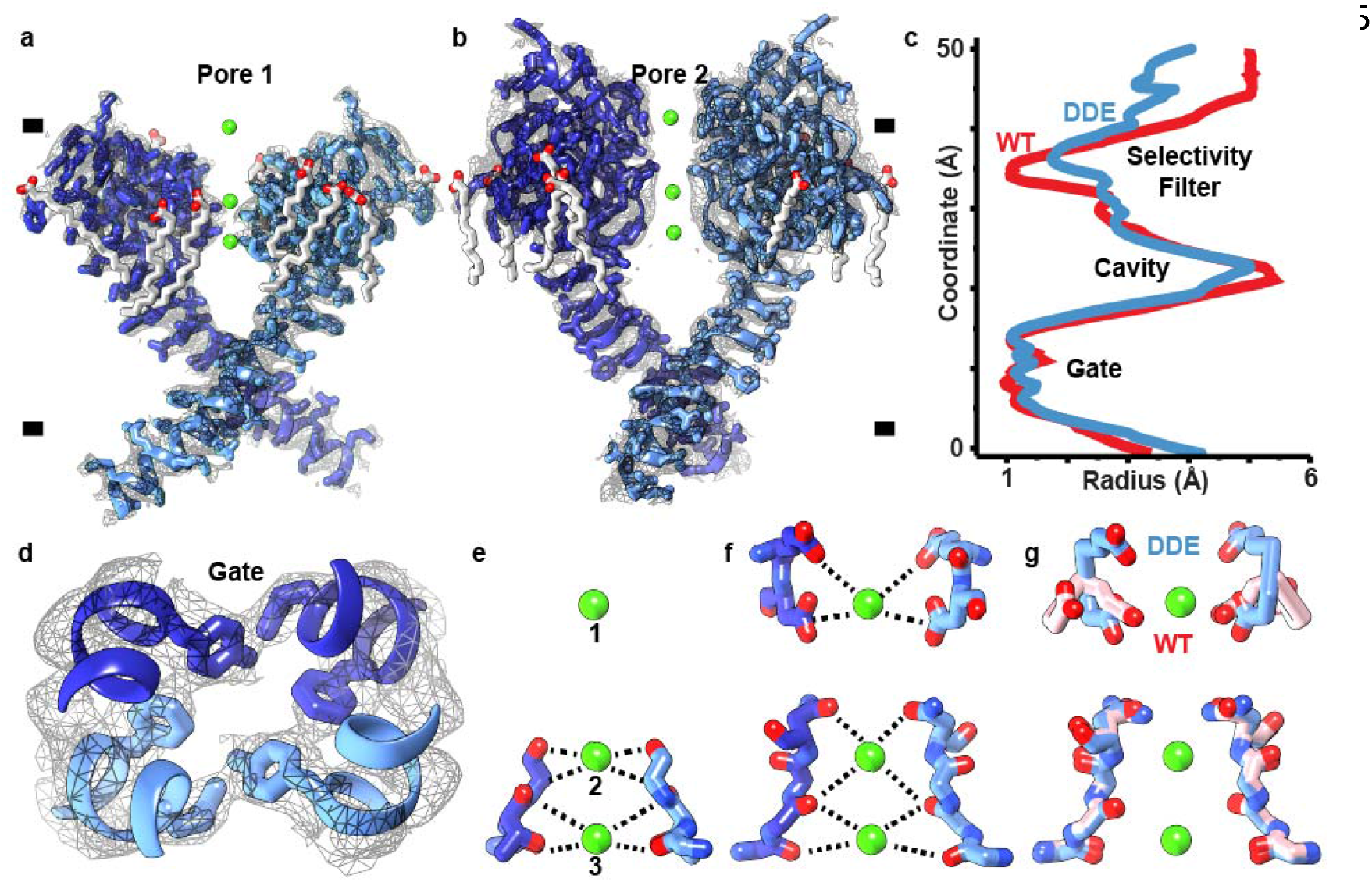
Ion Permeation Pathway. a, b, Orthogonal side views through pore helices (a) S5-S6 (pore 1) and (b) S11-S12 (pore 2) of the channel homo-dimer overlaid with high-resolution cryo-EM density (gray mesh). c, HOLE plot of pore radii along central channel coordinate of AtTPC1_DDE_ (red) and AtTPC1_WT_ (blue). d, Top down view through central pore. Gate residues Y305, L673, and F676 are shown. e-g, Side views through the selectivity filter in (e) pore 1, (f) pore 2, and (g) an overlay of pore 2 of AtTPC1_DDE_ (blue) and AtTPC1_WT_ (pink). Upper E605, D606 and lower S265, T263, T264, V628, M629, N631 selectivity filter residues are shown. Density for lipids (See Figure S7) and ions omitted for clarity.

In the selectivity filter, three ions are observed (Figure 4e, f). Site 1 and 3 were occupied by Ca^2+^ mimetics, Yb^3+^ or Ba^2+^, in AtTPC1_WT_ structures (19, 20). Site 2 was seen to be occupied by Na^+^ in the Na^+^-selective chimera of AtTPC1(33). Since AtTPC1_DDE_ contains 1 mM Ca^2+^ and the channel is Ca^2+^-selective, these are probably hydrated Ca^2+^-ions (See Supplemental Discussion). Sidechain interactions from the upper vestibule of S11-S12 exclusively coordinate site 1. The side chains of D606 move inward to coordinate site 1 with E605 (Figure 4g). E605-Ca^2+^ (5.5 Å) and D606-Ca^2+^ (4.6 Å) distances are consistent with the radius of a hydrated Ca^2+^ ion. There are readjustments throughout the channel suggesting that VSD2 activation by removal of the luminal Ca_i_^+^ site can indeed potentiate channel opening as observed by electrophysiology.

The observation of upper and lower selectivity filter and activation gate operating independently in our structures suggests a multi-step gating mechanism that resembles that proposed for TRPV1 channel activation (39). Activation of the VSDs could lead to opening of the selectivity filter and activation gate in two steps. First, the upper selectivity filter opens in response to rotational rearrangement and partial activation of the VSDs, then the lower selectivity filter and activation gate open when the VSD achieves maximal activation.

### Cytoplasmic activation by Ca^2+^-ions

Full voltage-dependent activation of AtTPC1 requires ~0.3 mM cytoplasmic Ca^2+^ evoked by Ca^2+^-binding to the EF3-EF4 loop (20, 29). D376 in EF3 is critical; when substituted D376A in AtTPC1_DA_ the channel remains closed and is no longer responsive to membrane potential (29). We determined the crystal structure of AtTPC1_DA_ (Figure 5a, Table S2) to 3.5 Å resolution by X-ray crystallography. When compared with the AtTPC1_WT_ crystal structure, removing the activating cytoplasmic Ca_a_^2+^ binding site leads to higher dynamic motion (B-factors) not only in the EF-hand and the CTD, but also in the VSDs and upper vestibule of the pore, showing that the activating cytoplasmic site may act through effects on the VSDs (See Supplemental Discussion).

**Figure 5.**
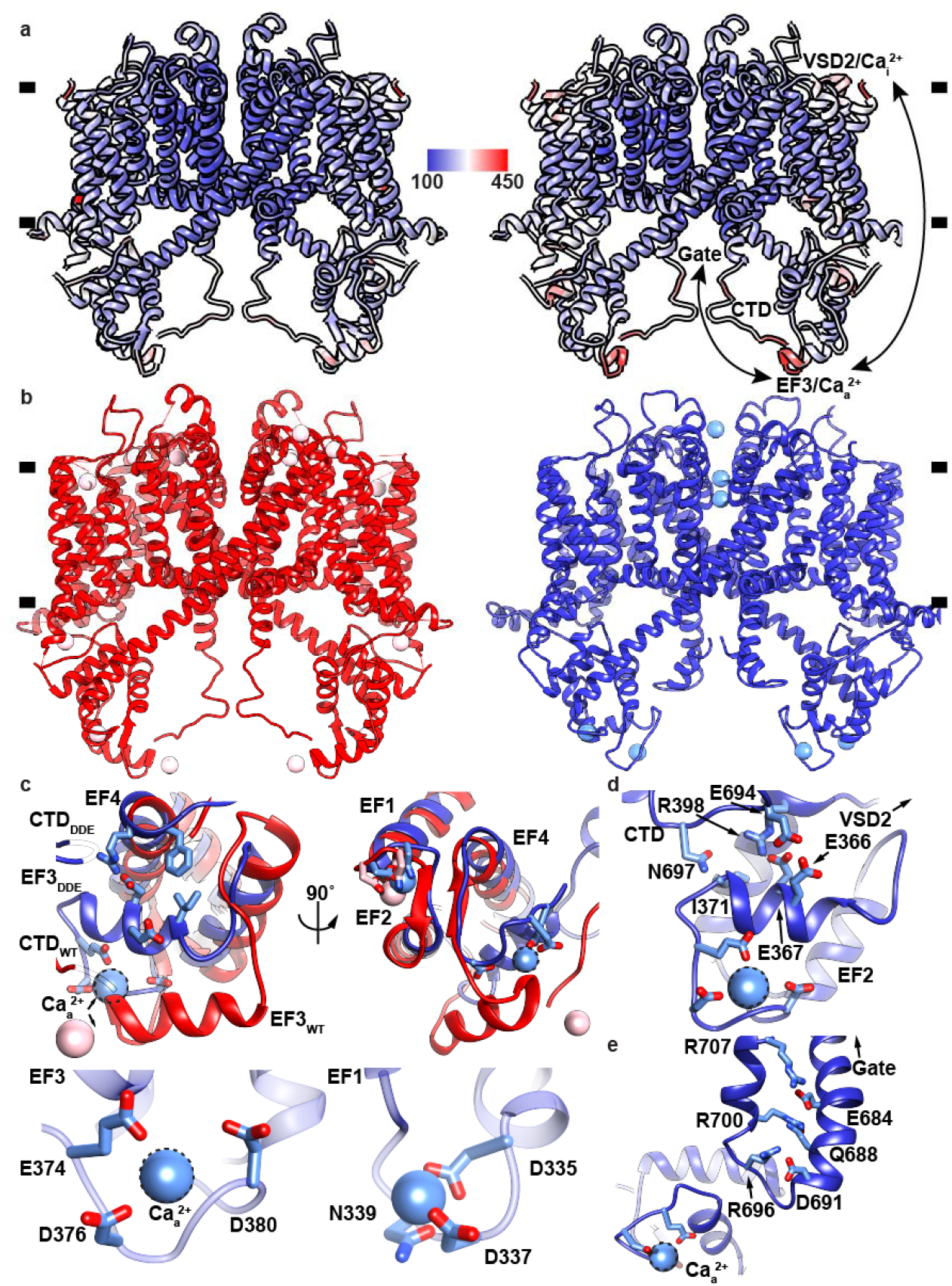
Dynamics of Cytoplasmic Domains. a, Overlay of (left) AtTPC1_WT_ and (right) AtTPC1_DA_ crystal structures colored by B-factor value (100-450 Å^2^) (See Methods and Supplemental Discussion). b, Sideviews of AtTPC1_WT_ (red) and AtTPC1 DDE (blue). c, Views from (top left) side and (top right) bottom of EF3 for overlaid AtTPC1_DDE_ (blue) and AtTPC1_WT_ crystal structures (red, PDBID 5DQQ). CTD interactions omitted for clarity. Bottom panels show Ca^2+^-ion binding sites in (bottom right) EF1-EF2 and (bottom left) EF3-EF4. d, View of EF3-CTD and EF3-EF4 interactions and connection to VSD2. e, Structure of the CTD in AtTPC1_DDE_ and a possible coupling pathway to the pore gate and Ca_a_^2+^. Dash lines indicate the hypothetical Ca_a_^2+^ ion position.

AtTPC1_DA_ is in a resting-state as expected. However, there is an overall effect of increasing the dynamical motion of VSD1, VSD2 and the pore (in the context of otherwise identical overall B-factors), showing that Ca_a_^2+^ has an allosteric effect in stabilization that could impact voltage dependence (VSDs) and conductance (the pore). The regions of increased motion in AtTPC1_DA_ correspond to the regions that undergo conformational change in AtTPC1_DDE_, suggesting that there is a pathway for coupling cytoplasmic activation at Caa^2+^ to voltage dependence of activation in VSD2 and transmission to the pore.

The Ca_a_^2+^ site in EF3 is fully formed in the cryo-EM structure of AtTPC1_DD_E, whereas it was partially occupied in AtTPC1_WT_ and was in a more extended conformation(19, 20) (Figure 5b). In AtTPC1_DDE_, EF3 alone lies 7 Å. closer to the transmembrane domain and rotates 20° to make close contacts with EF4 (Figure 5c). The Ca_a_^2+^chelating residues E374, D376, and D380 order the EF3-EF4 loop around the activating Ca_a_^2+^ site (Figure 5c). Movement of EF3 leaves the EF1-EF2 Ca^2+^ site unaltered. Therefore, EF3 is capable of undergoing large-scale movements that change the structure of the cytoplasmic domains and their connection to VSD2, suggesting that activation on the cytoplasmic side can act reciprocally through VSD2.

The conformation of the cytoplasmic domains seen in AtTPC1_DDE_, probably reflects the predominant structure in AtTPC1_WT_ (rather than the crystal structure) because the conformation of EF3 is conserved in cryo-EM structures from several conditions; detergent and saposin A nanoparticles of AtTPC1_WT_, and AtTPC1_DDE_, and when bound to either of two different Fab molecules (Figure S8c).

The CTD of AtTPC1 is indispensable for channel activation(40). Removal or truncation by 29 residues abolishes channel function. In the crystal structure of AtTPC1_WT_, the CTD forms an intramolecular complex with the EF3 via salt-bridge interaction D376-R700, leading to the hypothesis that the EF3-CTD complex could undergo conformational changes upon activating cytoplasmic Ca^2+^-binding (19). In the cryo-EM structures the CTD moves toward the membrane vertically by 13Å to accommodate the upward movement of EF3 upon forming the cytoplasmic activating Ca_a_^2+^ site (Figure 5b). The salt-bridge (D376-R700) that previously linked the CTD to EF3 in AtTPC1 _WT_ breaks to allow D376 to chelate the Ca_a_^2+^ site. This can explain why only substitution of D376 and not the other chelating residues abolishes Ca^2+^-activation(29). In AtTPC1_DDE_, the CTD now forms a hydrogen bond (E366-E694) with EF3 and hydrogen bonds (N697-I371) via the backbone (Figure 5d). Following the interaction with EF3, and a β-turn the CTD to folds back on itself to form a charged zippered interaction (R696-D691, R700-Q688, R707-E684) between the signal poly-R and poly-E motifs of the CTD (19). The poly-E motif connects directly to the pore gate (Figure 5e). The role of the CTD on channel activation may be to stabilize the EF3 helix in both apo- and Ca^2+^-bound conformations to allow communication of EF3 movement to VSD2 and the pore gate. Upon removal of Ca^2+^, the CTD could adopt an extended conformation to reform the salt-bridge interaction D376-R700 with EF3. Without the CTD, EF3 may not be able to reform the Ca_a_^2+^ site and would become trapped in an inactive state.

To further investigate the conformation of EF3 in solution, we performed continuous wave electron paramagnetic resonance (CW-EPR) experiments using spin-labeled full-length AtTPC1 lacking cysteines (AtTPC1_cysless_; See Methods) in detergent micelles. Ten positions in the CTD, gate, and EF-hand domains were examined by spin-probe mobility and responsiveness to Ca^2+^-ions. Labeling at a site on EF3 (R379) indicates a conformational shift to higher probe mobility as compared with EGTA, indicating that the probe changes environment in the presence of Ca^2+^-ions (Figure S9). Mutation of the Ca_a_^2+^ site but not the Ca^2+^-site in EF1-EF2 abolishes the Ca^2+^-dependent increase in probe mobility, suggesting that EF3/Ca_a_^2+^ changes conformation upon increasing cytoplasmic Ca^2+^-ion concentrations. The data suggest that the AtTPC1_WT_ crystal structure likely represents an apo-state of EF3, whereas AtTPC1_DDE_ represents a Ca^2+^-bound conformation present along the pathway of activation.

### Alternate Access Mechanism of Activation

As the electric potential changes across a membrane, functional voltage sensors find a new free energy minimum. Under hyperpolarizing conditions, the favorable electrostatic free energy component of positive gating charges’ attraction to internal negative potential is balanced against a structural ‘distortion’ free energy cost. Depolarization therefore releases both of these components to populate a new overall equilibrium state that includes a component that regulates the open probability of the pore. Release of the electrostatic free energy is achieved by alternating access of the positively charged gating charges on S10 to solvent from the cytoplasmic side in the hyperpolarized resting state, to a state in which they are insulated from the cytoplasmic side, and become accessible to solvent from the lumenal side, without any necessity for vertical movement of the gating charges themselves. The structures of AtTPC1_WT_ and AtTPC1_DDE_ suggest that this ‘alternating access’ mechanism plays the key role. The movement of the VSD2 helices S7, S8, S9 with respect to the gating charges on S10 achieve this (Figure 6).

**Figure 6.**
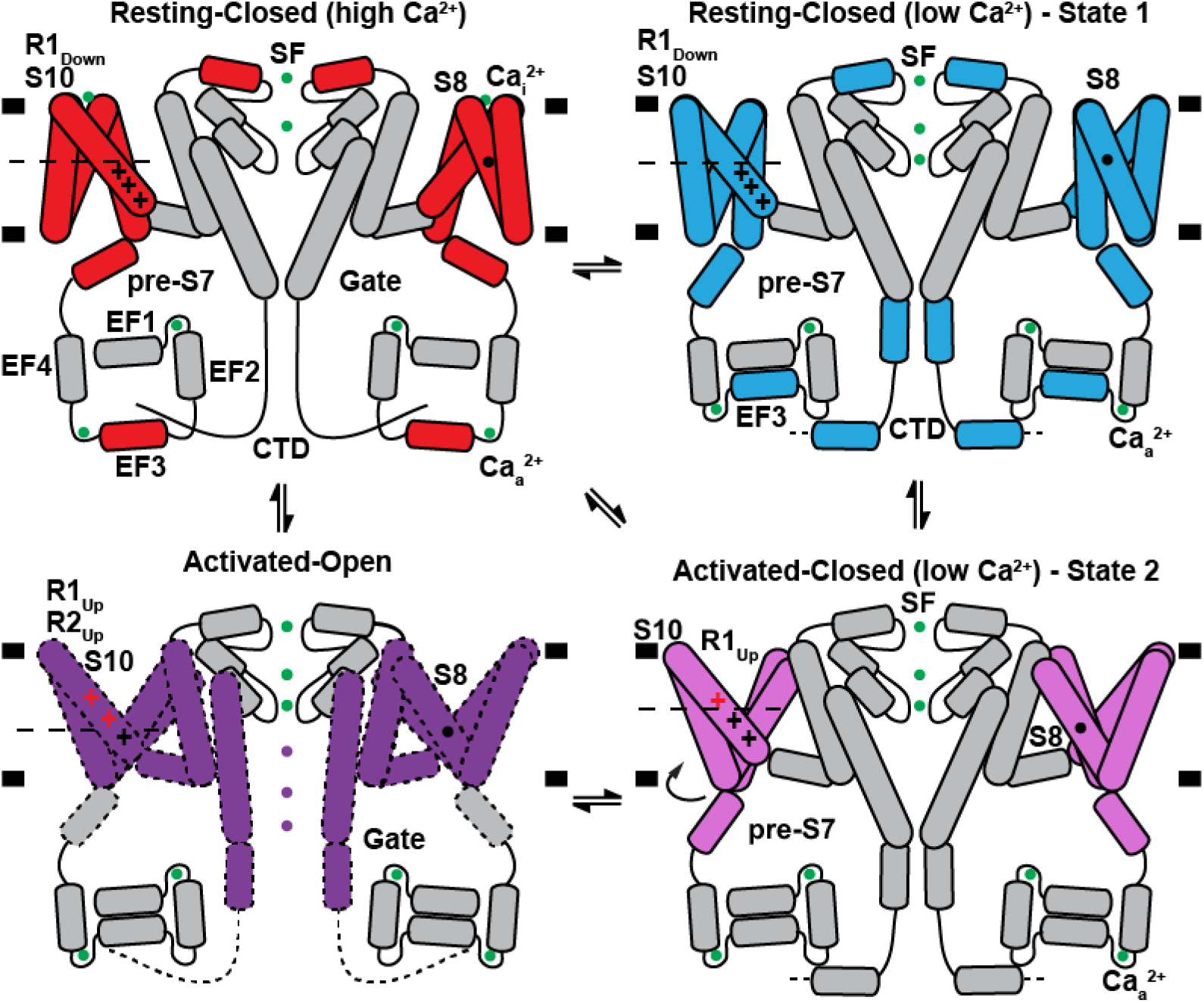
Mechanism of TPC1 Channel Activation. Schematic summarizing the conformations of VSD2, EF3, CTD, and selectivity filter (SF) observed in crystal and cryo-EM structures under high and low effective lumenal Ca^2+^-ion concentrations. A potential model for coupling between lumenal inhibition Ca_i_^2+^, cytoplasmic activation Ca_a_^2+^, voltage-sensing, and channel gating is shown. Helices and ions that move between the AtTPC1_WT_ resting-state (red), AtTPC1_DDE_ state 1 (cyan), AtTPC1_DDE_ state 2 (pink), and hypothetical activated-open state (purple) are colored. Ca^2+^-ions are shown as green balls. Gating charges that change position with respect to the CT (dashed line) between states are colored red.

Hence the potential gradient is focused across a thin hydrophobic region close to the lumenal (external) side in the resting-state, and alternately across one on the cytoplasmic side in the activated-state. Thus, the gating charges do not require any vertical movement across the bilayer as the electrically insulating region, which includes the CT and across which the voltage gradient is formed, moves from outside of the gating charges in the resting-state voltage sensor, to the cytoplasmic side of gating charges in the activated-state. Thus, the electric component of free energy is released.

The mechanism is unique versus most previously proposed models in that S7-S9, not only S10 that is closest to the pore, are crucial during voltage-activation to move gating charges from the electric potential on one side of the membrane to the potential of the other side without any necessary vertical translation of S10. Even after this major driving effect, gating charge side chains will also readjust in context of their different environments and some vertical movement may find more favorable energetics in the shallow energy gradient around them.

There is a teleologically favorable aspect to this alternate access mechanism in that it does not need to remove any hydrophobic residues out of the bilayer as would be required of a model involving large vertical movement of S10. It also does not need to rebuild the hydrophobic interface between S10 of the VSD and the walls of the pore with which it forms the primary contact that may be important in changes that alter open probability of the channel. Simulations of VSDs that assume a linear gradient of electric potential across the bilayer that changes only in slope on depolarization, but not position of start and end points across the VSD may not represent the true physiological state.

The Gibbs free energy change on moving one charge across a transmembrane potential difference of −70 mV is −1.6 kcal/mole. The measured gating charge transfer for activation of AtTPC1 is 3.9 charges(20) or −6.3 kcal/mole in free energy difference. Ca^2+^ that effectively stabilizes the resting-state. The EC_50_ for the Ca_i_^2+^ site is ~0.1 mM. Therefore, assuming a K_d_~0.1 mM, the free energy released by removing each Ca_i_^2+^ site is ~5.6 kcal/mole per site (or ~11.2 kcal/mole per AtTPC1 channel). It is not unreasonable that this difference, in releasing the restraint at Ca_i_^2+^, could allow the VSD to reach its active state that would normally be evoked by depolarizing the transmembrane potential at low Ca^2+^. The fact that we observe intermediate states of TPC1 upon removing Ca_i_^2+^ rather than the fully activated state suggests that the membrane voltage may contribute additional energy to maximize activation of VSD2 and channel open probability.

The mechanism suggested for AtTPC1 voltage dependent activation is the first time that atomic structures have been determined for intermediates on transition from resting-state to activated-state in an intact voltage-dependent channel. The structures also capture large-scale rotation of VSDs that could play a role in activation. It could be a general way in which gating charge can move energetically downhill in response to a change in voltage, without the energetic cost of removing any hydrophobic lipophilic regions out of the membrane.

## Acknowledgements

We thank J. Finer-Moore for aiding with manual building, refinement, and critical reading of the manuscript. We thank A. Brilot, K. Verba, E. Palovcak, and D. Asarnow for help with data processing and refinement, R. Wang for help using Rosetta, D. Bulkley and M. Braunfeld at UCSF, and Z. Yu and his colleagues at the HHMI Janelia Cryo-EM Facility for aiding with data collection, and Ali Punjani for a pre-release version of cryoSPARC v2. This work was supported by NIH grant GM24485 (to R.M.S), Postdoctoral Independent Research Grant (to A.F.K) from the University of California, San Francisco Program for Breakthrough Biomedical Research, which is partially funded by the Sandler Foundation, NIH grants R01GM098672, S10OD020054, and S10OD021741 (to Y.C.), NIH grants GM117372 and GM087519 (to A.A.K), and NIH grants R01EY05216 and P30EY00331, and the Jules Stein Professor Endowment, and the Bruce Ford and Anne Smith Bundy Foundation (to W.H.). E.M.G was supported by the National Science Foundation Graduate Research Fellowship NSF 1144247. Beamline 8.3.1 at the Advanced Light Source is operated by the University of California Office of the President, Multicampus Research Programs and Initiatives grant MR-15-328599. The Berkeley Center for Structural Biology is supported in part by NIH, NIGMS, and HHMI. The Advanced Light Source is supported Contract No. DE-AC02-05CH11231. Stanford Synchrotron Radiation Lightsource Contract No. DE-AC02-76SF00515. The SSRL Structural Molecular Biology Program is supported by the DOE, and P41GM103393. Chimera was supported by NIGMS P41-GM103311.

## Author Contributions

A.F.K. conceived and designed the project with assistance of R.M.S. A.F.K. and P.K.D. purified TPC1. A.F.K. performed crystallization and structure determination experiments, refined crystal and cryo-EM structures, and optimized saposin A reconstitution; E.M.G. collected and processed cryo-EM data with J-P.A. and Y.C.; P.K.D. reconstituted TPC1 into nanodiscs and generated Fabs with D.D., S.S.K., and A.A.K.; M.B. collected EPR data with W.H.; A.F.K., E.M.G., Y.C., and R.M.S. wrote the manuscript. All authors contributed to manuscript preparation.

## Methods

### Protein Production

#### AtTPC1

X-ray and cryo-EM trials used AtTPC1 with deletion of residues 2-11(19). Antibody generation employed full-length AtTPC1. AtTPC1 constructs were expressed and purified by nickel affinity chromatography (NiNTA), thrombin cleavage, and size exclusion, as described previously(19). For reconstitution into saposin A lipoprotein nanoparticles and MSP nanodiscs, the size exclusion step was omitted after NiNTA, but performed after reconstitution. To introduce single cysteine labels for CW-EPR, a cysteine-less variant of AtTPC1, AtTPC1_cysless_, was constructed to replace all native cysteines with serines (C93S, C101S, C159S, C347S, C392S, C574S, C577S, C580S, C687S, C728S). Cysteine substitutions for EPR spectroscopy were purified in the presence of 1mM TCEP during solubilization and binding to Nickel beads, but then removed during wash and elution steps to prevent interference with MTSL-labeling. AtTPC1_cysless_ has identical biochemical behavior to wild-type AtTPC1.

#### Saposin A

Human saposin A (plasmid was a kind gift from J. Frauenfeld) was expressed and purified from *E. coli* Rosetta-gami 2(DE3) cells essentially as described(30). Several colonies from transformed cells were used to inoculate overnight 300 mL LB cultures containing 25 μg/mL chloramphenicol, 10 μg/mL tetracycline, and 15 μg/mL kanamycin at 37 °C. Overnight cultures were used to seed 6 L of Terrific Broth, grown to an absorbance value at 600 nm of 1. The temperature was adjusted to 20 °C 30 minutes prior to induction by 0.7 mM IPTG for 15 hours at 20 °C. Cells were harvested by centrifugation at 4,000x g for 10 minutes and resuspended in 200 mL 50 mM tris pH 7.4, 150 mM NaCl (Lysis Buffer) and frozen at −80 °C before further use. Bacterial pellets were lysed in the presence of 1 mM phenylmethylsulfonyl fluoride (PMSF) by sonication for 5 minutes, then clarified by centrifugation at 26,000x g for 20 minutes. The supernatant was then incubated at 70 °C for 10 minutes followed by centrifugation at 26,000x g for 20 minutes to remove contaminant proteins. Imidazole pH 7.4 was then added to 20 mM and passed through two HisTrap FF crude 5mL columns equilibrated in Lysis Buffer with 20 mM Imidazole pH 7.4 using a peristaltic pump. Following 50 mL washes with Lysis Buffer with 20 mM Imidazole pH 7.4 and Lysis Buffer with 35 mM Imidazole pH 7.4, pure human saposin A was eluted in 25 mL Lysis Buffer with 400 mM Imidazole pH 7.4 in 1.5 mL fractions. Fractions containing human saposin A were dialyzed against 1 L of Lysis Buffer with 4 mg of Tobacco Etch Virus (TEV) protease. The following morning an additional 2 mg of TEV was added with an exchange of the dialysis buffer and dialyzed at room temperature for 3 hours. Uncleaved saposin A and free TEV were removed by passing the protein through a single HisTrap FF crude 5 mL column equilibrated in Lysis Buffer with 20 mM Imidazole pH 7.4 and washed with 30 mL of Lysis Buffer with 20 mM Imidazole pH 7.4. The flow-through and wash fractions were pooled and concentrated in 5 kDa molecular weight cutoff concentrators and serially injected in 20 mg aliquots over TSKgel G3000SW 7 mm x 60 cm or Superdex S200 10/300 columns equilibrated in Lysis Buffer with 1 mM CaCl_2_. High and low molecular weight peaks were observed during size exclusion. The low molecular weight, most consistent with the size of the saposin A monomers and dimers, was concentrated to 2 mg/mL for use in reconstitution of AtTPC1 and frozen at −80 °C. Final yields of human saposin A were 5-10 milligrams of human saposin A per liter of culture.

#### Membrane scaffold protein

Plasmid harboring His-tagged E3D1 variant of membrane scaffold protein (MSP) of the nanodiscs was obtained from Addgene (#20066). MSP was expressed, purified and cleaved with TEV protease as described(41) with minor modifications. Colonies of transformed BL21 (DE3) Gold cells (Stratagene) were used to inoculate overnight 30 mL LB cultures containing 30 μg/mL kanamycin. Overnight cultures were used to inoculate 2 L of Terrific Broth, grown at 37 °C until OD_600_ of 1.3 and then induced with 1 mM IPTG for 3 hrs. Cultures were harvested by centrifugation at 4,000x g for 10 minutes, resuspended in 50 mM Tris pH 8.0, 300 mM NaCl (MSP Lysis Buffer) supplemented with 1% Triton X-100, protease inhibitor tablet and 1 mM PMSF. Microtip sonicator (Branson) was used to lyse the cells for 3 cycles at 1s on, 1 off and at 60% amplitude for 3 minutes each. For each 1L culture 1 mL of NiNTA resin equilibrated with MSP Lysis Buffer was used in batch-binding for 1 hour at 4 °C. Resin was then washed with MSP Lysis Buffer supplemented with: 1) 1% Triton X-100 - 10 column volumes (CVs), 2) 50 mM sodium cholate, 20 mM imidazole – 5 CVs, 3) 50 mM imidazole – 7 CVs. MSP was eluted with 4 CVs of MSP Lysis Buffer supplemented with 400 mM imidazole. TEV cleavage was performed overnight in dialysis against 4 L of 50 mM Hepes pH 7.5 and 200 mM NaCl. Uncut protein was removed with subtractive step on NiNTA resin and cleaved MSP was concentrated to 5 mg/mL using Millipore concentrators with 10 kDa molecular weight cutoff and frozen in aliquots at −80 °C. Typical yield of MSP E3D1 from 1 L culture was 15 mg.

#### Antibodies

Monoclonal antibodies were expressed and purified from hybridoma culture supernatants using standard methods in the Monoclonal Antibody Core Facility at OHSU (Dan Cawley). 4B8 Fab fragment was generated by papain digestion 1/20 Papain:4B8 (w/w) at 30 °C for 1 hour in the presence of 5 mM L-cysteine and quenched with 20 mM iodoacetamide. Full-length IgG and Fc regions were removed via protein A chromatography and the flow-through containing 4B8 Fab was concentrated and purified by TSK3000/Superdex200 size exclusion chromatography in 20 mM Hepes pH 7.4, 0.2M NaCl, 5% Glycerol.

Antibody fragments (Fabs) generated by phage display were expressed and purified essentially as described(42) from constructs subcloned into the expression vectors pRH2.2 or pSFV4 (gift from S. Sidhu). DNA was transformed into BL21-Gold(DE3) (Agilent) and used directly to set up 15 mL overnight starter cultures in 2xYT media supplemented with 50 ug/mL ampicillin. Cultures were then inoculated into 1 L of 2xYT media, grown until OD_600_ of ~0.8-1.0, induced with 1 mM IPTG for 4 hours at 37 °C. Cells were harvested by centrifugation and disrupted in lysis buffer containing 20 mM sodium phosphate dibasic pH 7.4, 500 mM NaCl, 1 mM PMSF, and 2 mM DNase, using sonicator (Branson). Lysate was heated to 55 °C for 30 minutes, cleared by centrifugation and incubated with 1 mL of MabSelect ProteinA resin (GE Healthcare) in batch for 1 hour at 4 °C. Resin was washed with 15 column volumes of 20 mM sodium phosphate dibasic pH 7.4, 500 mM NaCl and then eluted with 10 ml 0.1 M acetic acid into tubes containing 1 mL of 1 M Hepes pH 7.5 to immediately neutralize elutions containing Fab fragments. Fabs were then dialyzed against 4 L of 50 mM HEPES pH 7.5 and 200 mM NaCl, concentrated with Millipore concentrators with 30 kDa molecular weight cutoff and stored in aliquots at −80 °C. Typical yield from 1 L culture varied depending on Fab fragment and was between 0.5 – 3 mg. For cryo-EM studies Fab CAT06 was modified with H12 elbow variant as described previously(43), resulting in the CAT06/H12 antibody fragment. Briefly, the H12 elbow variant exchanges heavy chain residues (112)SSASTK(117) (numbering according to Kabat (44)) to (112)FNQI-K(117) with one position deleted from the sequence.

### Saposin A and Nanodisc Reconstitution

#### Lipids

All lipids were prepared from thin films. Lipids were dissolved in chloroform, aliquoted into glass vials, dried under nitrogen, and stored at −20 °C dry. Unless otherwise noted, resuspending lipids was accomplished by adding buffer with or without detergent and sonicating 10-30 minutes until clear.

#### Saposin A nanoparticles

Saposin A nanoparticles containing AtTPC1 were formed by mixing AtTPC1 alone or AtTPC1-Cat06/H12 Fab complexes with soy polar lipids and saposin A in a 1:8:5-10 ratio by mass. A typical preparation consisted of 2.4 mg (8 mg/mL) of NiNTA purified AtTPC1 incubated with 2.4 mg (9 mg/mL) of protein A purified CAT06/H12 (adjusted to 0.05% DDM, 1 mM CaCl_2_), incubated at room temperature for 5-10 minutes. AtTPC1-Cat06/H12 Fab complexes were then mixed with 4.8 mL of 5 mg/mL soy polar lipids (diluted from a 25 mg/mL stock in 1% DDM) dissolved in Size Exclusion Buffer (20 mM Hepes pH 7.3, 0.2 M NaCl, 5% Glycerol, 1 mM CaCl_2_, 0.05% DDM) and incubated at room temperature for 30 minutes. Then 9.6 mL (2 mg/mL) saposin A was added, incubated 1 hour at room temperature, then diluted to 40 mL with Size Exclusion Buffer. Detergent was removed by adding 1 g of activated, washed, Bio-Beads SM2 (BIO-RAD) resin overnight at 4°C. Bio-Beads were activated by incubation in methanol and successively washed with copious amounts of water. The following morning, beads were removed by filtration through 5 μm syringe filter. 0.5 molar equivalent of Fab CAT06/H12 was added, incubated 30 minutes at 4 °C, then concentrated to ~0.3 mL in a 100 kDa MWCO concentrator, and filtered prior to injection on a TSK4000 or Superose 6 column equilibrated in Nanodisc Size Exclusion Buffer (20 mM Hepes pH 7.3, 0.2 M NaCl, 1 mM CaCl2). A single fraction was selected for cryo-EM analysis by screening for homogeneous particles by negative-stain EM (Figure S6b-d). Cryo-EM grids were frozen following ultracentrifugation at 100,000x g (TLA-55) for 5 minutes at 4 °C without further concentration. AtTPC1-CAT06/H12-saposin A complexes typically eluted from size exclusion at 0.5-1 mg/mL. 16 lipids are observed bound to AtTPC1_DDE_ (Figure S7).

#### MSP nanodiscs

Reconstitution of AtTPC1 into E3D1 nanodiscs was performed using standard protocols(45). For the purpose of antibody generation by phage display, full-length AtTPC1_WT_ was reconstituted into nanodiscs using biotinylated version of MSP E3D1, and size exclusion fractions corresponding to AtTPC1 in biotinylated nanodiscs were used without the removal of empty nanodiscs. Final mixed lipid-detergent micelles used in reconstitution contained 10 mM soy polar extract, 1 mM CHS, and 30 mM DDM. Nanodisc assembly mix was prepared using purified TPC1, mixed micelles and MSP at 2:10:1000 ratio. All components were mixed together in Nanodisc Size Exclusion Buffer and incubated on ice for 1 hour. Next, 400-500 mg activated Bio-Beads were added per each ml of assembly mix, and the reconstitution reaction was left shaking overnight at 4°C. As a control, empty nanodiscs were prepared using the same phospholipids. All nanodisc samples for phage display experiments were concentrated, supplemented with 5% w/v sucrose, aliquoted, flash-frozen in liquid nitrogen, and kept at −80°C.

#### Proteoliposomes

Proteoliposomes of full-length wild-type AtTPC1 were formed by resuspending 20mg of soy polar lipids dried from chloroform resuspended in 4 mL Nanodisc Size Exclusion Buffer, subjected to 10x alternating liquid Nitrogen and warm water bath freeze-thaw cycles, then sonicated for 10 min. The opaque mixture was then passed through an Avestin hand extruder with a 400 nm pore size 10x until mildly translucent. Vesicles were disrupted by addition of 69.8 μL 10% DDM added (4 mM final), 12.1 μL 2% CHS in 10% DDM added (0.1 mM final), and 1 mM CaCl_2_, and stirred 30 minutes at room temperature. 2 mg of AtTPC1 was added to disrupted liposomes (1:10 protein:lipid ratio) and stirred for 1 hour at room temperature using a magnetic “flea” stir bar. To form proteoliposomes, methanol washed Bio-Beads were added in increments of 300 mg, 300 mg, 500 mg, and 1 g in 1-hour intervals at room temperature. The last incubation was done at 4 °C overnight. The following morning the samples were centrifuged at 100,000x g for 1 hour in a table-top ultracentrifuge (TLA-55). The proteoliposome pellet was resuspended in in 0.4 mL of Nanodisc Size Exclusion Buffer. The concentration was estimated by gel and frozen in liquid Nitrogen and stored at −80 °C. Empty liposomes were made in a similar fashion by substituting AtTPC1 protein with Size Exclusion Buffer.

### Antibody Generation

#### Hybridoma

Hybridoma cell lines were generated by standard methods at OHSU by Daniel Cawley. Briefly, four Balb/c mice were injected with 25 μg of AtTPC1 proteoliposomes at 0 and 14 days. On day 26, serum was prepared and tested for antibody titer in ELISA. The mice with the highest titers were used to derive hybridomas. The two best responding mice were injected on day 45-55 with 10 μg Ag as a final boost. 4 days later, spleen cells are fused with P3X mouse myeloma cells. Hybridomas were assayed 12 days later in ELISA. All candidate cell lines were expanded and given a secondary ELISA screen 12-14 days later. All antigen specific antibodies were made available as culture supernates for screening. All candidate cell lines are frozen in liquid nitrogen. 14 hybridoma antibodies were identified that specifically bind to immobilized biotinylated AtTPC1 in DDM detergent by ELISA. To select for antibodies that recognize 3D structural epitopes a counter-selection against antibodies that bind linear epitopes on denatured protein in 8M Urea was performed, yielding 5 antibodies that bind only under native conditions by ELISA. Selection of high-affinity antibodies that were capable of complex formation was conducted by fluorescence size exclusion chromatography(46) (FSEC) using FITC-labeled AtTPC1 and hybridoma supernatants containing 10-50 μg/mL whole IgG. FSEC samples were prepared by mixing 1μg FITC-Labeled AtTPC1 with 0.4 mL hybridoma supernatants (supplemented with components to make Size exclusion buffer +0.03 mg/mL Soy Polar Lipids) in a total volume of 490 μL of Size exclusion buffer. Samples were incubated at 25 °C for 1 hour or 4 °C for 4-5 hours and injected on a Superose 6 column, 0.4 mL/min, equilibrated in Size exclusion buffer at 4 °C, using an autosampler and inline fluorescence detector set at excitation=495 nm and emission=518 nm. The fluorescence detector (Shimadzu RF-10AXL) was calibrated before use by injecting varying amounts of FITC-AtTPC1 (0.1 ng-1 μg) with known labelling efficiency.

#### Phage Display

To generate Fabs, the efficiency of biotinylation of AtTPC1_WT_-E3D1 nanodiscs was evaluated by pull-down on streptavidin-coated magnetic beads (Promega). Library sorting steps were performed using Fab Library E (DNA kindly provided by S. Koide(47)) based on previous protocols(31, 48). Six independent phage library sorting experiments were performed against biotinylated AtTPC1_WT_ -E3D1 nanodiscs in 2 buffers containing 20 mM Hepes pH 7.3, 200 mM NaCl, 5% glycerol, 1% BSA, and either 1 mM CaCl_2_ or 1 mM EGTA. Additionally, for each CaCl_2_ or EGTA condition, sorting was carried out in the presence of following ligands: either 1 μM NAADP, 1 μM trans-NED19, or no ligand control to maximize the number of multiple states of TPC1 channel during sorting (Sorting Buffer). Total of five rounds of phage library sorting were performed for each sample with decreasing concentration of immobilized biotinylated TPC1-E3D1 nanodiscs between subsequent rounds in following order 1000 nM, 600 nM, 200 nM, 200 nM and 100 nM. Round 1 of sorting was performed manually using nanodisc bound to streptavidin-coated magnetic beads, and upon washing with respective Sorting Buffer, whole beads were used to infect log phase *E. coli* XL-1 strain (Stratagene) in the 2xYT media (Fisher) supplemented with 50 μg/ml ampicillin and 10^9^ pfu/ml of KO7 helper phage (NEB) overnight at 37 °C and 280 rpm to amplify the phage particles. The amplified phage particles after round 1 were used as an input for four additional rounds of library sorting performed semi-automatically with a KingFisher magnetic bead handler (Thermo) according to described protocols(49). Starting from round 2, in every step except elution, the buffer was supplemented with 10-fold molar excess of non-biotinylated empty E3D1 nanodiscs to counter-select for MSP-, lipid-, and non-specific phage particles. Finally, in each of rounds 2-5, phage particles were eluted from magnetic beads by 15-minutes incubation with 1% Fos-Choline-12 in respective Sorting Buffer.

Initial validation of selection clones was performed by single point direct phage ELISA using clones from round 3, 4 and 5. Amplified phage particles at 10-fold dilution were assayed against 50 nM biotinylated nanodiscs (either empty or with AtTPC1) using HRP-conjugated anti-M13 monoclonal antibody (GE Healthcare, #27-9421-01). Assays were performed in respective Sorting Buffer supplemented with 2% BSA. Each Fab clone with A_450_ signal above 0.2 (four times the average background level of the assay) was sequenced; unique Fabs were sub-cloned into pSVF4 or pRH2.2 vectors (kind gift of S. Sidhu), and purified as described above. In total 16 unique Fab sequences were obtained from total of 192 single colonies screened; 9 Fabs from library sorting with CaCl_2_ and 7 Fabs from library sorting with EGTA.

### Cryo-EM Structure Determination

#### Sample Preparation

Samples were analyzed by negative-stain EM to determine suitability for cryo-EM. To prepare grids, 3 μl of sample at 10–50 μg ml^−1^ was applied to a glow discharged continuous carbon grid, which was then treated with 0.75% (w/v) uranyl formate. Grids were imaged on an FEI Tecnai T12 microscope operated at 120kV at a nominal magnification of 52,000× using an UltraScan 4000 camera (Gatan) or an FEI Tecnai T20 microscope operated at 200kV and at a nominal magnification of 80,000× using a TemCam-F816 8k × 8k CMOS camera (TVIPS) camera, corresponding to pixel sizes of 2.21 Å and 0.95 Å, respectively, on the specimen.

Grids for cryo-EM were prepared with FEI Mark IV vitrobot. Quantafoil R1.2/1.3 400 mesh holey carbon grids (EMS) were glow-discharged for 30 s. Then 2.5 μl protein sample at a concentration of 0.6-1.5 mg ml^−1^ was applied onto the carbon face of the grids. The grids were blotted with Whatman#1 filter paper for 2-4 s at 100% humidity and plunge frozen in liquid ethane. The grids were loaded onto a 300kV Polara (FEI) with a K2 Summit direct electron detector (Gatan). Data were collected at nominal magnification of 31,000×, corresponding to a physical pixel size of 1.2156 Å (0.6078 Å super resolution pixel size) on the specimen with a dose rate of 7.6 electrons per physical pixel per second. Images were recorded with SerialEM in super-resolution counting mode and a defocus range of −0.8 to −2.0 μm. A total exposure of 12 s was used, with 0.2 s subframes (60 total frames) to give a total dose of 60 electrons per Å^2^ (1.35 electrons per Å^2^ per subframe). Data for AtTPC1_WT_ in saposin A was also collected at the Janelia Cryo-EM facility FEI Titan Krios microscope operated at 300kV with a K2 camera with a physical pixel size of 1.02 Å.

#### Image Processing

For negative-stain data, GCTF(50), Gautomatch (Kai Zhang, url: https://www.mrc-lmb.cam.ac.uk/kzhang/), and RELION-2(51) were used for CTF estimation, particle picking, and 2D classification, respectively. For cryo-EM data, dose-fractionated super-resolution image stacks were drift corrected and binned 2×2 by Fourier cropping using MotionCor2(52) (after discarding the first frame). CTF determination and particle picking was performed on motion-corrected sums without dose-weighting using GCTF and Gautomatch. To generate particle picking templates and initial models a Gaussian reference was used to pick particles and 2D classification was performed in RELION-2 followed by *ab-initio* reconstruction in cryoSPARC. Particles were then picked using six 2D classes as templates. 2D classification, 3D classification and refinement procedures were carried out using and RELION-2 and cryoSPARC(53) (Figure S3). After 3D classification the reconstruction was filtered to 30 Å resolution and used as an initial reference model for 3D refinement in cryoSPARC v2 using the beta version of non-uniform refinement yielding a reconstruction with a gold-standard Fourier shell correlation of 3.5 Å with a 0.143 cutoff criterion. The map from cryoSPARC was used for modeling the Fab variable domain. The final 3.3 Å high-resolution reconstruction was performed in RELION-2 using 3D auto-refinement by masking out the saposin A membrane belt and the Fab using a mask created from a 30 Å low-pass filtered model of AtTPC1_WT_. Local resolution estimates were performed using RELION-2. The final maps were sharpened in RELION-2 using a B-factor of −104 and −117 Å^2^ for the TPC1-Fab-saposin and TPC1-only reconstructions, respectively.

State 1 and 2 were identified by focused classification of VSD2 without image alignment using the angles determined from the high-resolution reconstruction (Figure S3). Focused classification benefitted from applying C2 symmetry. After 3D classification each particle set was refined in RELION-2 with global angular searches using a 30Å low pass filtered reconstruction of the combined dataset. The maps for state 1 and 2 extend to 3.7 Å as determined by the gold-standard FSC. These maps were used to build atomic models for VSD2 with local resolution in this region ranging from 4-6 Å as estimated by RELION-2 (Table S1). The final maps were sharpened in RELION-2 using a B-factor of −95 and −86 Å^2^ for the state and state 2 reconstructions, respectively.

#### Structure Determination and Refinement

The entire structure of AtTPC1_DDE_ excluding VSD2 was built into the highest resolution map (High-res; Table S1) manually after initial real-space flexible fitting and refinement of the AtTPC1_WT_ structure into the map using Rosetta (54) with the electron scattering table. Manual fitting in COOT(55), followed by real-space coordinate and B-factor/atomic displacement factor refinement in PHENIX(56) were used in *de novo* building of the NTD, EF-hand, CTD, Ca^2+^-ions, lipid molecules (palmitic acid and phosphatidic acid), and the Fab-AtTPC1_DDE_ interface. This map was calculated with a mask around TPC1, excluding the Fab. A structure of the Fab-complex (Fab-bound; Table S1) was determined from the same dataset by extending the mask to include the Fab variable domains (V_H_ and V_L_). The Fab was built using a homologous structure of a Fab from the same library (PDB 3PGF) with the variable CDR loops deleted.

Building of the state 1 (State 1; Table S1)was accomplished by rigid body alignment of the AtTPC1_WT_ structure to the cryo-EM map using CHIMERA (57) followed by flexible fitting and refinement in Rosetta, and extensive manual fitting in COOT. State 2 (State 2; Table S1) was built by first rotating VSD2 of AtTPC1_WT_ to place S10. Movement of S7-S9 around the S10 axis was done manually in real-space. Gating charges R1-R3 were placed in favored rotamer positions using the rotamer library in COOT and PHENIX, while applying the additional restraint that R1-R3 must contact solvent, a polar side chain, or counter charge in the membrane. One position satisfied these criteria for each gating charge.

Model validation employed MolProbity(58) and EMRinger(59). Low resolution maps of AtTPC1_WT_-4B8 were interpreted by flexible fitting of the AtTPC1_WT_ crystal structure and a homology model of 4B8 in real-space using Rosetta. Final AtTPC1_DDE_ coordinates and maps have been deposited in the Electron Microscopy Data Bank (EMDB) under EMDB IDs (High-resolution) 8957, (Fab) 8956, (State 1) 8958, and (State 2) 8960, and the Protein Data Bank (PDB) under PDB IDs 6E1M, 6E1K, 6E1N, and 6E1P.

#### Crystal Structure Determination

AtTPC1 D376A (AtTPC1_DA_) crystals were obtained in the presence of CaCl_2_, CHS, soy polar lipids, and 1mM trans-NED19 (NED19), using HiLiDe(60) as described previously(19). Average diffraction was 4 Å, with 10% diffracting to 3.5-3.7 Å resolution, the best being 3.5 Å. Anisotropic resolution was determined using the CCP4 program Truncate(61) and the UCLA Anisotropy Server(62). Crystals were partially dehydrated by incubation with additional 15% (v/v) of polyethylene glycol 300 prior to freezing in liquid nitrogen.

X-ray diffraction datasets were collected at the Advanced Light Source (ALS) Beamlines 8.3.1 and 5.0.2 and at Stanford Synchrotron Radiation Lightsource (SSRL) Beamline 12-2. Data were reduced using XDS(63). The best native dataset extends to overall resolution 3.5 x 6.0 x 4.5 Å (ref.(62)). Phases were calculated by molecular replacement using PHASER(64) and AtTPC1 wild-type (PDBID 5DQQ) as a search model. Structure interpretation was using COOT(55), with refinement in PHENIX(56). The structure of AtTPC1_DA_ was refined to 3.5 Å resolution with final R_work_/R_free_ of 31.84 % and 35.19%. A sharpening B-factor of −142.05 Å^2^ was used for refinement, as described previously(19). Analysis by Molprobity shows Ramachandran geometries of 93.09%, 6.25%, and 0.66% for favored, allowed, and outliers. The structure contains 11.31% rotamer outliers. For comparison of B-factors between AtTPC1_WT_ (mean Wilson B-factor 108 Å^2^; PDB ID 5DQQ)(19) and AtTPC1DA (mean Wilson B-factor 109.86 Å^2^) structures, B-factors were first scaled by adding the difference in mean wilson B-factor (1.86 Å^2^) to the mean wilson B-factor of the wild-type AtTPC1 crystal structure (PDBID 5DQQ)(19). The AtTPC1_DA_ structure and structure factors have been deposited in the Protein Data Bank (PDB) under PDB ID 6CX0.

### EPR Spectroscopy

#### Sample Preparation for CW-EPR

Concentrated (7-12 mg/mL) NiNTA-purified AtTPC1_cysless_ mutations were incubated with 10-fold molar excess MTSL (MTSL; Toronto Research Chemicals, Inc) dissolved at 10 mg/mL in DMSO for 2 hours at room temperature. The samples were then split in half and treated with either 1mM CaCl_2_ or 5mM EGTA pH 7.4 prior to filtration and purification by size exclusion chromatography in 1mM CaCl_2_ or 5mM EGTA pH 7.4, respectively. AtTPC1_cysless_ has identical biochemical behavior to AtTPC1_WT_ and maintains structural integrity as evidenced by negative-stain EM analysis. Fractions containing labeled AtTPC1_cysless_ were pooled and concentrated to 5 mg/mL for EPR measurements.

#### EPR Data Collection

For continuous wave (CW) EPR experiments, X-band spectra were collected on a Varian E-102 Century series spectrometer fitted with a loop-gap resonator (Medical Advances, Milwaukee, WI). Samples (10 uL) were contained in a 0.64/0.80 mm (i.d./o.d.) quartz capillary (Vitrocom, Mountain Lakes, NJ) and spectra were recorded at room temperature over 100 G with an incident microwave power of 2 mW, modulation amplitude of 1 G, and modulation frequency of 100 kHz; typical total scan times were 5 minutes. CW spectra were normalized and corrected for minute deviations in baseline prior to comparison and analysis. All data were plotted using Origin 6 after normalizing the area under integrated CW spectrum.

Cryo-EM structural data of AtTPC1_DDE_ has been deposited to the Protein Data Bank under accession codes 6E1K, 6E1M, 6E1N, 6E1P and the Electron Microscopy Data Bank under accession codes 8956, 8957, 8958, 8960. X-ray structural data of AtTPC1_DA_ has been deposited to the Protein Data Bank under accession code 6CX0. Correspondence and requests for materials should be addressed to R.M.S and Y.C.

